# Cryo-EM protein structure without purification

**DOI:** 10.1101/2025.08.13.669967

**Authors:** Samantha M. Powell, James E. Evans

**Affiliations:** Pacific Northwest National Laboratory, Environmental Molecular Sciences Laboratory, Richland, WA, USA; Washington State University Pullman, School of Biological Sciences, Pullman, WA, USA

**Keywords:** Cryo-EM, Cell-free Expression, Lysate, Protein Structure

## Abstract

Cryo-electron microscopy (cryo-EM) is a powerful technique for macromolecular structure determination. While several aspects of the workflow are either semi- or fully automated, the beginning of the workflow related to the generation of macromolecular samples remains a bottleneck limiting full automation. In most cases, the production of the initial sample still takes days to months. Initial sample production often takes days to months. Here, we show that cryo-EM can solve structures of proteins directly from cell-free expression complex mixtures with minimal or no purification while also reducing reagent volumes and overall labor and time. Using a cell-free reaction volume of just 25 microliters, we achieve rapid protein synthesis and structural characterization in under 24 hours at resolutions better than 3Å. These results highlight a lower cost and rapid gene to structure workflow with minimal perturbation satisfies the foundational requirements needed to facilitate future improvements in throughput via full workflow automation.

## Introduction

Protein structure determination is fundamental for understanding the functions and mechanisms of biological macromolecules since the three-dimensional (3D) arrangement of atoms in a protein dictates its interactions, stability, enzymatic activity, and binding properties. Historically, determining protein structures has been a laborious and time-consuming process, often taking years to complete^1^. However, the last 5 years have witnessed increasing demand for rapid protein structure determination especially following the release of AlphaFold ^2^ and other computational approaches that permit in silico and de novo structure prediction in hours to days. While these computational approaches have high accuracy for many proteins, challenging protein groups still have low accuracy and require experimental validation. Improving the throughput for high-resolution structural characterization of such proteins under near-native buffer/environmental conditions is needed to keep pace with the field demands.

Cryo electron microscopy (cryo-EM) had a resolution revolution a decade ago ^3^ and was the focus of the Nobel Prize for Chemistry in 2017 ^4^. It has exploded in recent years as a structural biology approach capable of reaching atomic resolution and that does not require protein crystallization. The typical workflow ^1^ for cryo-EM single particle analysis (SPA) involves four major components: 1) generating the sample, 2) vitrifying and optimizing sample distribution on cryo-EM grids, 3) high-resolution single particle data collection, and 4) image processing which encompasses particle curation, 2D/3D refinement and model building/validation. Modern electron microscopes can now handle and screen 12 cryo-EM grids automatically. In conjunction with automated data collection and real-time image processing software that permit real-time, and in some cases user-free, 3D refinement ^5^, the desire to improve throughput to match rates compatible with sampling 96 or more samples in one day on one microscope has driven researchers toward developing automated sample handling and grid vitrification as well as faster direct electron detectors. This in turn has created a need for faster and more cost-effective methods for generating the target proteins in a manner compatible with cryo-EM and downstream automation simply to keep pace with future instrument capacity.

Generating the sample has always represented the first major bottleneck for the cryo-EM workflow. Most commonly, protein purification is included as a part of sample preparation, but it’s a time-consuming and often challenging step. To combat this, many groups have utilized affinity grids using a variety of different tags, antibodies, and/or grid substrates to allow cell lysates to be applied to a TEM grid for on-grid purification.^6-10^ Unfortunately, these approaches often result in further challenges, such as the accessibility to the tag, the specificity of binding, or the presence of the tag hindering native folding or assembly of the target protein. Other groups have explored visualizing over-expressed cell lysates by directly lysing the cells after adsorption to a TEM grid substrate, but the surrounding debris can complicate analysis. Recently, the ‘Build-and-Retrieve’ (BaR) method was introduced, where a lysed and fractionated sample that is partially cleaned up with size exclusion chromatography is deposited directly on a grid for determination of multiple structures from the mixed population. ^11-15^ While the BaR method permits analyzing heterogeneous samples and has the benefit of working with native complexes, the scale of the sample harvesting and laborious purification steps is still too large for our purpose toward full automation. We aim to synthesize only 1-5 micrograms of the target protein which is the projected amount of protein needed for combined small scale biochemical analysis, functional analysis, and structure determination from a single sample.

Cell-free expression pipelines have previously been used to generate proteins in a test tube for structural and functional annotation^16,17^ but have generally been more focused on toxic proteins or complex heterocomplexes that are difficult to express and purify from cell-based methods. While protein synthesis in a test tube for direct structural analysis has permitted going from gene to high-resolution cryo-EM protein structure in less than two weeks after receiving a custom synthesized gene clone^18^, that example remains an outlier, and the conventional workflow still takes a few months on average for most proteins. Here, we present a method for structure determination directly from a miniaturized (25 µl) cell-free expression reaction with zero, or minimal purification. The power of our method is that it minimizes reagents, time, and labor associated with generating the protein target, and it reduces and/or eliminates purification steps needed for sample preparation for cryo-EM single particle analysis. The ability to use untagged proteins also eliminates many of the above challenges and can increase the likelihood of determining the true native structure of a protein. Furthermore, although the protein expression system being used in our approach generates a final sample that is heterogeneous, all components are well-defined and known and can be computationally identified and separately processed to allow the entire process from expression to structure determination to occur in less than 24 hours.

## Results

Single particle cryo-electron microscopy (cryo-EM) is a leading technique for high-resolution 3D structure determination of macromolecular complexes. Despite its transformative impact on structural biology, several inherent limitations of cryo-EM become particularly pronounced when dealing with complex mixtures. First and foremost, the presence of heterogeneity within the sample is usually considered undesirable ^12^. Single particle analysis approaches rely on generating structures from 2D projections of “equivalent” particles at random orientations. This means the traditional paradigm for cryo-EM is to use samples with maximum homogeneity to increase likelihood of successful structure determination ^19^. Meanwhile, the presence of heterogeneity can lead to blurred or ambiguous density maps if the averaging process employed by cryo-EM struggles to reconcile multiple structural states ^20^. 3D classification and variability analysis algorithms exist but the iterative refinement can be computationally expensive. Ideally, we would like to utilize an approach that allows rapid 2D classification of the various proteins of most abundance within a given sample. Since our method would intentionally utilize complex mixtures, we needed to ensure that we identified the best cell-free expression lysate and system that would permit high yield expression, minimize total reagent requirements, minimize the timeframe for expression and has an overall component list that wouldn’t interfere with effective particle picking or lead to misidentification or misclassification of particles as these could introduce systematic errors in the reconstruction process.

### Choosing the appropriate cell-free lysate

We previously developed a cell-free expression pipeline based upon wheat germ lysates and have used a robot synthesizer to perform coupled expression and affinity purification at the 1.2 ml or 6 ml scale for cryo-EM structure determination of the target proteins. Using this, the timescale from receiving a clone and completing wheat germ-based cell-free expression, purification and structure determination at 2.5Å was 9 days ^18^. We have also previously shown that 55 microliter scale reactions of the wheat germ cell-free expression system can be used to test for expression yield, solubility, and native folding and assembly of the target protein by supplementing with pre-charged FluoroTect Green lysine tRNA (Promega, WI, USA), and that 220 microliter scale reactions are compatible with direct functional assays using the crude expression mixture. While the yields and automation compatibility are favorable for this wheat germ system, we ran a proteomic analysis of the reaction lysate and identified ∼3000 unique proteins present in the lysate even before synthesizing the protein target of interest. Such a large number of components was deemed to be untenable for the current approach since the typical yield for the expressed target proteins did not fall within the top 20 protein components in the mixture. Thus, we expected that the likelihood of detecting proteins in cryo-EM micrographs without purification would be low, therefore we sought a different source expression system.

The PURE (protein synthesis using recombinant elements) system^21^ was originally published in 2001 and has since then been commercialized. The PURE cell-free expression system offers several notable benefits including a high protein yield, small scale reactions, and rapid setup and expression timeframes. However, the most important feature of this system for enabling the structure without purification workflow was its well-defined and minimal components. The PURE system ^22^ is known to be composed of only 36 total protein complexes (purified proteins, tRNAs, ribosome, and other necessary factors) and the typical target protein expression level places it within the top 5 components based on average abundance. This represents a 100x decrease of unique proteins that would be present in the background and that would complicate cryo-EM analysis. We therefore utilized the PURE system exclusively in all experiments reported here.

#### Artemia ferritin structure without purification as a benchmark

With the PURE cell-free system identified, we sought to address whether it could be used to rapidly produce protein in a test tube at a small scale and then solve the protein structure directly from the crude lysate. We settled on a 25-microliter scale for the PURE cell-free reactions as initial biochemical analysis showed that these were sufficient for detecting the target protein expression with both SDS and Native PAGE while leaving half the reaction available for dedication to cryo-EM grid preparation. As seen in Figure 1, we performed rapid cell-free expression by mixing a clone of ferritin from *Artemia franciscana* (Uniprot Q8WQM7) with reagents for in-vitro transcription and translation all in a 25 µl reaction and incubated at 37°C for 2 hours. This clone was chosen since this particular protein has not had a structure experimentally determined yet, but it shares homology with the conserved eukaryotic ferritin protein family that is widely used as a standard test sample with cryo-EM. The SDS and Native PAGE results (Figure 2 A&B) clearly depict the ferritin expression and assembly into its octahedral homomeric complex as expected. Direct deposition of the crude lysate to cryo-EM grids followed by grid vitrification, screening and data collection yielded micrographs with obvious ferritin particles (hollow sphere 2D projection characteristic of ferritin family members) surrounded by the complex milieu of the crude lysate (Figure 2 C).

**Figure 1.**
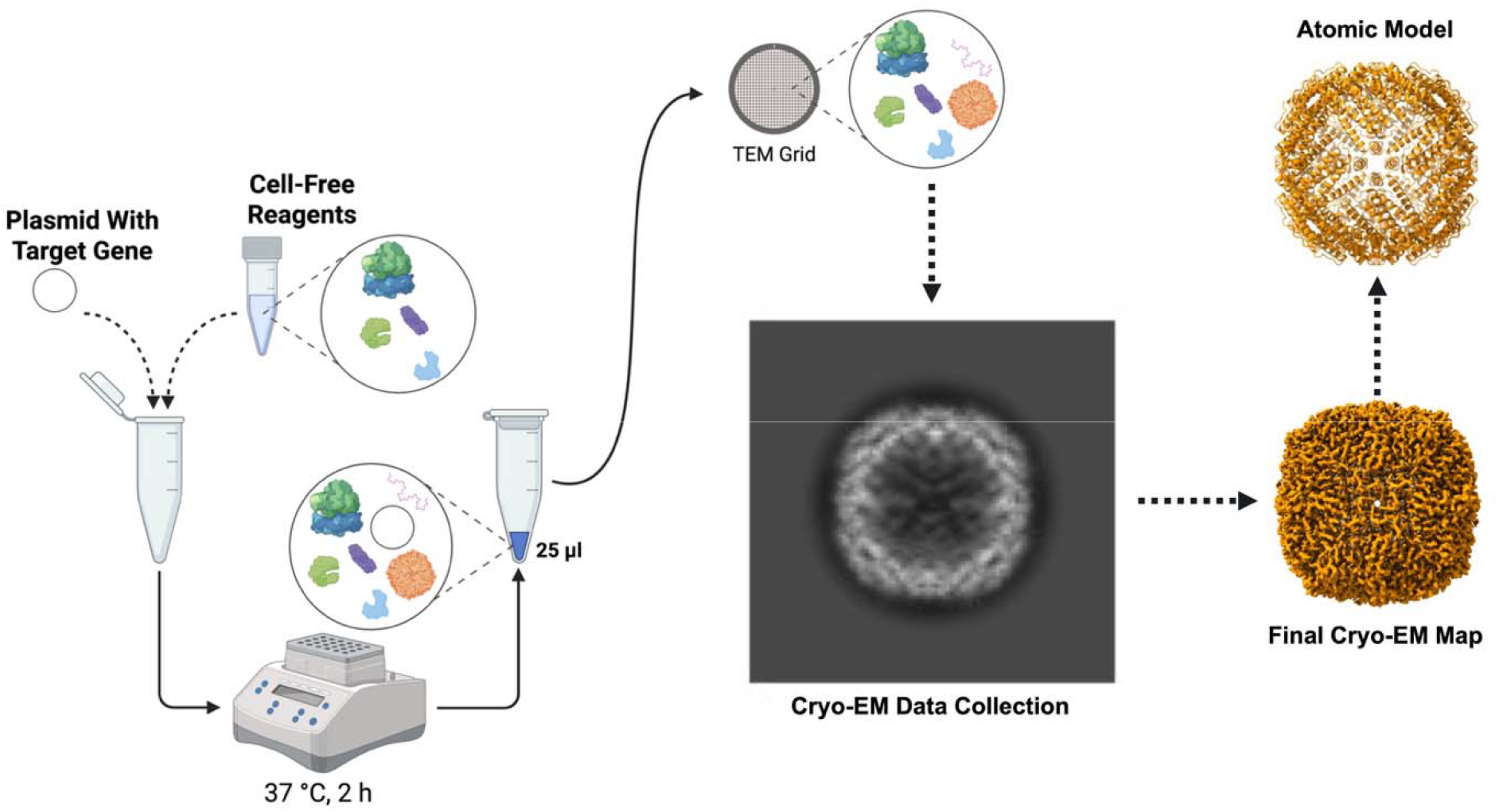
Overview of the structure without purification approach. Schematic of the structure without purification workflow from small scale cell-free expression reactions. Miniaturized protein production in a 25µl cell-free expression reaction provides ample target protein yields for suitable cryo-EM structure determination direct from lysate. A plasmid encoding for the target protein undergoes cell-free expression using the PURE *E. coli* based in-vitro transcription and translation in a test tube incubated for 2 hours. The resulting crude lysate containing all components of the cell-free lysate as well as the input plasmid DNA, transcribed mRNA and translated target protein is then loaded onto a cryo-TEM grid, vitrified and imaged with single particle cryo-EM. Following image processing a final cryo-EM derived map is generated that is used to fit and refine an atomic model of the target protein. Created in BioRender. Evans, J. (2025) https://BioRender.com/8lnvjsg

**Figure 2.**
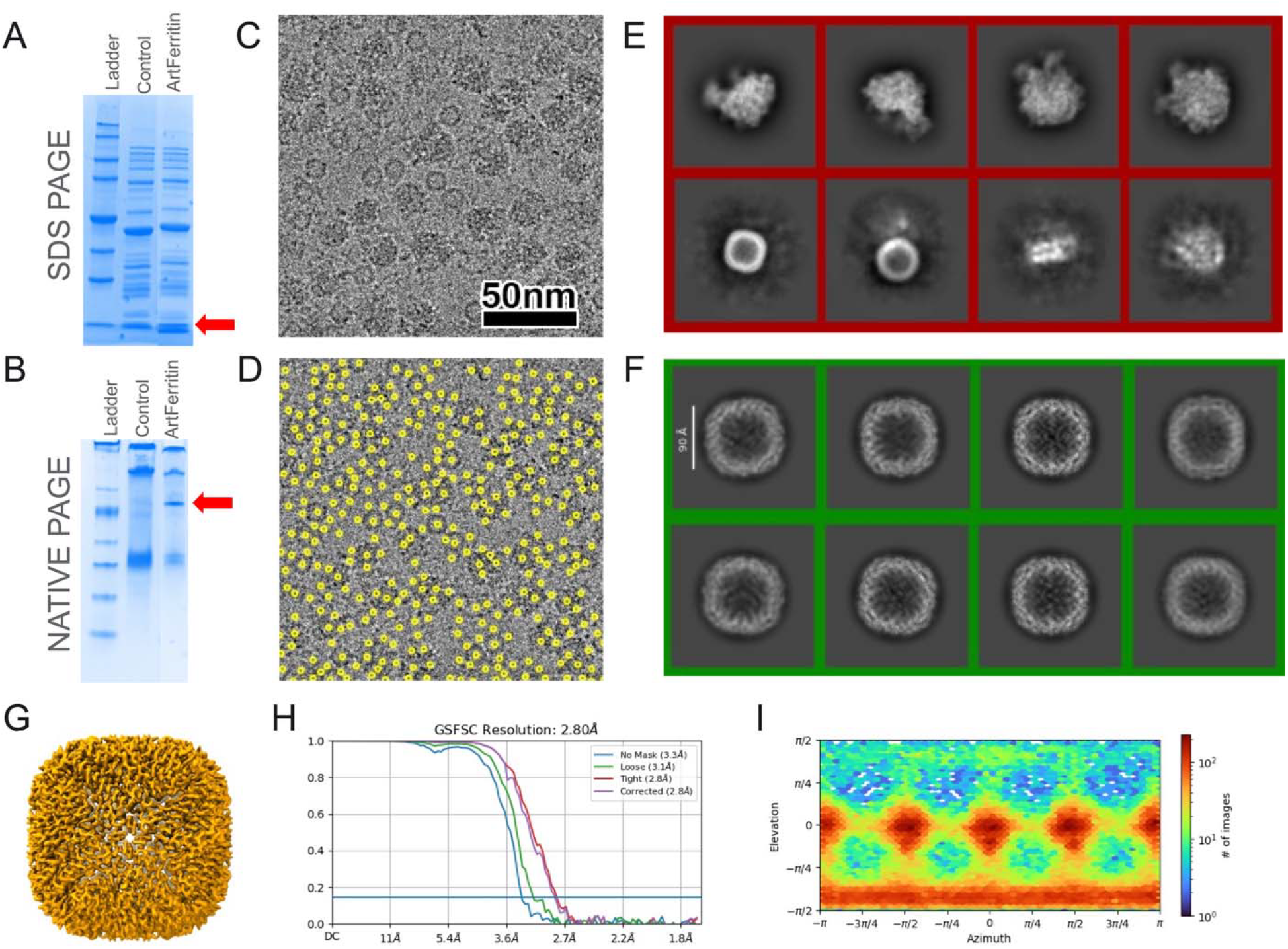
Ferritin “structure without purification” approach. A) SDS and (B) Native PAGE analysis of cell-free expressed ferritin compared to a no plasmid DNA control. The arrows point at unique bands indicating the expressed ferritin monomer in SDS PAGE and the assembled 24-mer in Native PAGE. C) Representative micrograph of ferritin complexes suspended in vitreous ice along with all other components from the lysate. D) Same micrograph from (C) with initial blob particle pick locations shown in yellow. E) Reference-free 2D class averages of picked particles showing the ability to discriminate classes clearly from the octahedral ferritin complex in the first round of 2D classification to identify and exclude ribosomes. F) A second round of 2D classification identifies the initial classes of our target protein. G) Surface rendering of final 3D refined map with corresponding Gold-Standard Fourier Shell Correlation plot (H), and viewing direction distribution plot (I) highlighting a good diversity of views present in the final dataset.

Cryo-electron microscopy micrographs were initially screened and manually inspected to ensure adequate coverage of particles amidst the complex background of cell-free expression lysate. The mixture contained target ferritin protein, plasmid DNA, transcribed mRNA, and the various components necessary for in-vitro transcription and translation including ribosomes. Using automated particle picking software, approximately 250,000 particles were identified (Figure 2 E). Initial 2D classification separated potential ferritin particles from noise and non-target elements within the lysate. We found that two rounds of 2D classification were sufficient for the initial particle triaging (Supplementary Figure 1). In the first round, all particle positions were boxed out with a total box size of ∼320Å and the 2D classification ran with a mask diameter of 270Å to identify ribosome particles (Figure 2E) that could be excluded from future steps. The non-ribosome classes were selected and used for the second round of classification using a mask diameter of 132Å to identify ferritin particles (Figure 2F). Subsequent iterations of reference-free 2D classification yielded clearer classes that represented distinct views of ferritin. From these, ∼20,000 particles were selected and used to generate an ab initio model that was then iteratively refined through several rounds of 3D classification and alignment to improve particle homogeneity and resolution. The final 3D reconstruction was generated from a total population of 90,000 particles and achieved an overall resolution of 2.8 Å (Figure 2 G-I), as determined by gold-standard Fourier shell correlation (FSC) at FSC=0.143 criterion. The resulting density map was of high quality, displaying well-resolved secondary structural elements such as the classical four alpha-helix bundle, with the short fifth helix at the C-terminus of each of the 24-subunits that make up the homomeric ferritin complex with octahedral symmetry.

#### Some rapid and small-scale purification steps improve resolution for ferritin

While our original goal was to avoid purification, we also wanted to evaluate whether rapid, small-scale purification steps could improve resolution. Figure 3 shows a workflow where instead of subjecting the crude lysate directly to a TEM grid, the 25 µl reaction was subjected to one of four small scale purification methods: 1) centrifugation using a 300 kDa MWCO spin filter to remove smaller proteins and mRNA, 2) doping with benzonase enzyme for 30 minutes to degrade mRNA or plasmid DNA, 3) reverse His-purification with magnetic Ni-NTA beads to remove His-tagged PURE components, or 4) combining methods (e.g., His-purification with 1 MDa MWCO spin filtering to remove His-tagged PURE proteins and ribosomes). Single-particle cryo-EM datasets for each condition showed all additional purification steps, which added at most 1 hour to the pipeline, improved resolution for ferritin, extending the 2.7Å map to as high as 2.1Å. Crude lysates with minimal purification resolved most features of an ideal purified sample, with lysate components like plasmid DNA showing no interaction artifacts or structural perturbations in the final reconstruction. Interestingly, benzonase treatment showed nearly the same improvement as removing all His-tagged components, making both approaches amendable to automation in future workflows. The final map at 2.1Å, purified with reverse His-affinity, enabled direct model fitting and refinement with Phenix against AlphaFold3’s initial structure. Supplemental Figure 2 and Tables 1 and 2 summarize the final map and reconstruction statistics. Overall, this 90,000-particle structure derived from cell-free lysates in less than 24 hours featured clear side chain densities, enabling accurate polypeptide assignment and achieving a map-to-model correlation of over 80%.

**Figure 3.**
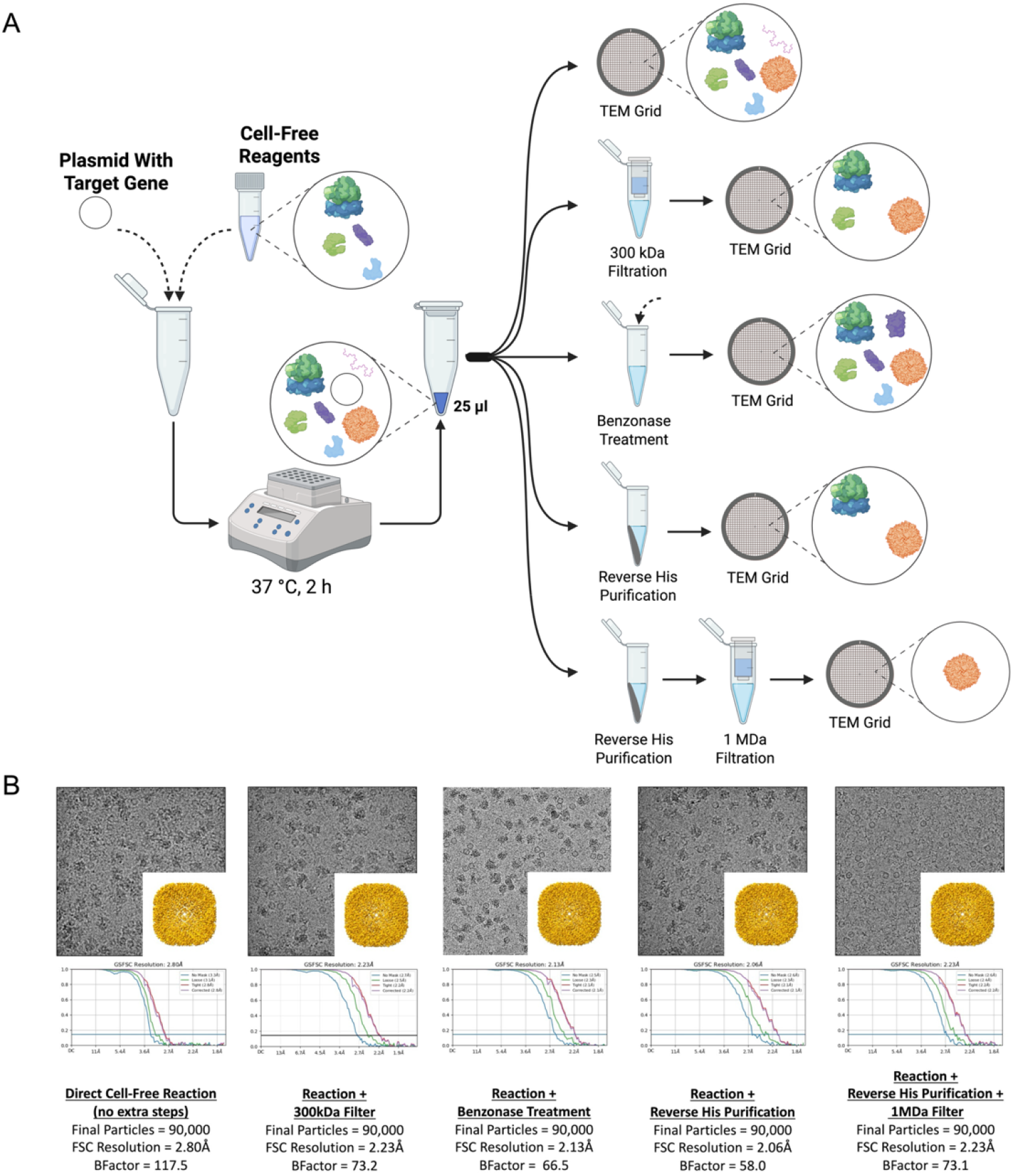
Comparison of attainable resolution for various post expression treatments. A) As seen in Fig. 1, a crude cell-free lysate can be directly imaged without purification. Alternatively additional small scale purification steps, that add <1 hour of extra sample handling and do not require more than the original 25 microliters as input, can be implemented to try and improve final resolution. Illustrated workflows for “cleaning up” the sample include small scale size exclusion filtration (imaging either filtrate or retentate to visualize complexes bigger or smaller than the cutoff), doping benzonase enzyme to degrade the mRNA and DNA present in the background, reverse His-purification where the majority of other protein components in the lysate are removed leaving only the untagged target protein and ribosomes in solution, beside the target protein, and finally multiple treatment such as combined reverse His-purification and spin column size exclusion filtration to remove all components other than the target protein. B) Comparative final structures from each of the workflows outlined in (A) using the same imaging parameters for all datasets including the number of particles in the final refinement. Note that simply doping in benzonase enzyme achieves approximately the same overall improvement as compared with the other approaches. Created in BioRender. Evans, J. (2025) https://BioRender.com/l0bp6e5

##### Benchmarking the structure without purification approach with a non-standard test sample of similar molecular weight as ferritin

Ferritin is a high-symmetry (24-mer) complex with molecular weight around 460 kDa. We aimed to test our new method on a similarly sized protein that wasn’t a standard cryo-EM test/calibration specimen and had less symmetry. Previously, we used large-scale cell-free expression and purification from wheat germ lysates to determine the structure of the *Arabidopsis* plant vitamin B6 biosynthesis protein PDX1.2 at 3.5 Å resolution (PDB 7LB5). This homomeric 12-mer (∼410 kDa) has roughly the same mass but half the symmetry of ferritin. Based on the ferritin results, we wondered if PDX1.2 would yield similar performance or worse resolution due to lower symmetry. Thus, we used PDX1.2 as a benchmark, as even purified PDX1.2 hasn’t been solved beyond 3Å with cryo-EM, although its x-ray structure is 2.0Å. The structure without purification method applied to PDX1.2 crude lysate (Figure 4) achieved 2.5 Å resolution, 1Å better than prior purified datasets, with the entire workflow taking only 18 hours from expression to refinement. Benzonase treatment or cleanup via small-scale purifications (Supplemental Figure 3/Supplemental Table 3) didn’t significantly improve resolution as it only extended to ∼2.4Å,. Fitting the known structure (PDB 7LB5) to the direct lysate map showed strong correlation and RMSD of <0.25Å compared to prior model coordinates.

**Figure 4.**
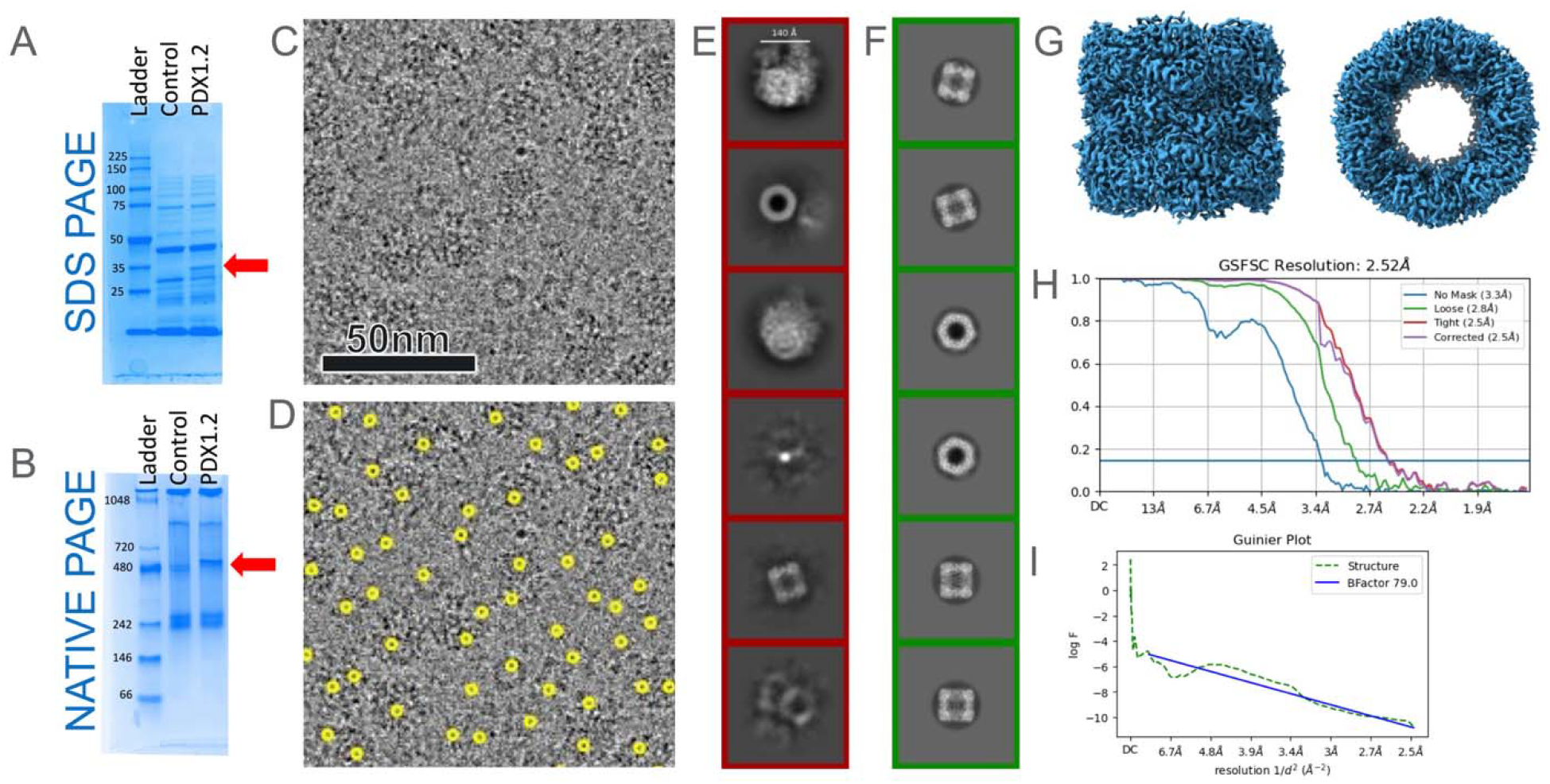
PDX1.2 structure without purification approach. (A) SDS and (B) Native PAGE analysis of cell-free expressed PDX1.2 compared to a no plasmid DNA control. The arrows point at unique bands indicating the expressed PDX1.2 monomer in SDS PAGE and the assembled 12-mer in Native PAGE. (C) Representative micrograph of PDX1.2 complexes suspended in vitreous ice along with all other components from the lysate. (D) Same micrograph from (C) with initial blob particle pick locations shown in red. (E) Reference-free 2D class averages of picked particles showing the ability to discriminate classes clearly from the dodecameric PDX1.2 complex (top 3 classes). (F-H) An overview of the cryoSPARC processing workflow showing (F) final 3D refined map, (G) Fourier Shell Correlation (FSC) curves, and (H) viewing direction distribution plot highlighting a good diversity of views present in the final dataset.

## Discussion

Our results demonstrate the successful application of a cell-free expression system combined with cryo-EM techniques for direct structural analysis of proteins without purification and using a minimal expression volume of only 25 microliters. We resolved the structure of a standard sample like ferritin to better than 3Å resolution in under 24 hours and achieved resolutions better than 2.4Å within 18 hours for a non-standard test protein. This streamlined approach opens new possibilities for rapid structural investigations of proteins, particularly those that are difficult to purify or toxic to cells. While the method achieves sub-3Å resolution, additional work is needed to test the limits of its applicability, such as compatibility with smaller protein complexes in terms of size and symmetry.

An unexpected but favorable discovery was that ribosomes present in vitrified samples stabilized vitrification conditions, enabling near-ideal ice formation and reducing the burden of grid screening. This finding supports automated grid preparation efforts, though further analysis is required to establish the range of proteins benefiting from ribosomes’ stabilizing effect and improving air-water interface interactions.

One common criticism of cell-free systems is that they lack full post-translational modification machinery or chaperones, raising concerns about proper folding or assembly. However, purified microsomes from multiple species are now commercially available, enabling reconstitution of active modification enzymes within cell-free workflows. Additionally, the PURE system’s simplified mechanics allow for genetic code expansion technologies, incorporating post-translational modifications through non-canonical amino acids. Furthermore, commercial options exist for functional chaperones (e.g., GroEL/ES, DNA J/K/E), enabling additive screens to optimize folding conditions for specific proteins.

In summary, we successfully demonstrated rapid protein production and structure determination using a streamlined workflow that combined efficient protein expression with cryo-EM. The PURE system accelerated sample production compared to complex in vivo systems, reducing costs and labor by shrinking the scale of expression experiments. By circumventing traditional bottlenecks in protein expression, we achieved sufficient yields for single-particle cryo-EM within hours and completed the workflow from sample production to structural refinement in less than 24 hours. This rapid protein synthesis approach holds enormous potential in structural biology, drug discovery, and functional studies, paving the way for high-throughput and fully automated sample screening.

## Online Methods and Materials

### Protein expression and characterization

Clones of *Artemia franciscana* ferritin (Uniprot Q8WQM7) and *Arabidopsis thaliana* Pyridoxal 5’-phosphate synthase-like subunit PDX1.2 (Uniprot Q9ZNR6) were acquired from Genscript using their custom gene synthesis and cloning services. The cloned genes were inserted into the pT7-DHFR Control plasmid (replacing the DHFR gene using NdeI and NotI restriction enzyme sites) provided in the PureExpress In Vitro Protein Synthesis kit (New England Biolabs) used for all cell-free expression experiments. All presented results utilized the 1x 25µL reaction scales per the standard protocol although the translation mixture was supplemented with FluoroTect GreenLys reagent (Promega) to permit fluorescent detection of the target proteins using SDS and Native PAGE.

For reverse His-tag based purification, a 1:1 ratio of expression reaction to packed magnetic Ni-NTA beads (HIS-Select Ni Magnetic Agarose Beads, Millipore) was used. The expression reaction was applied to the beads and incubated for 30 minutes, after which the purified sample was separated from the beads using a magnet. Benzonase treatment was carried out by incubating an expression reaction supplemented with 1 mM MgCl2, 2 mM CaCl2, and 5x diluted benzonase (Sigma) for 30 min at 30°C. All size exclusion filtration was performed using microcentrifuge spin filters (Sartorius Vivaspin 500 centrifugal concentrators) of various MWCO, centrifuged at 12,000xg.

### Cryo-EM sample preparation and single particle data collection

Three µL of each sample solution was loaded onto glow discharged Quantifoil 300 mesh R2/1. Grids were blotted for 0.5-1 s and plunge frozen in liquid ethane on a Leica EM GP2. Grids were stored in liquid nitrogen until further use. For screening and data collection, grids were loaded on a 300 keV Titan Krios G3i (Thermo Fisher) and all datasets were collected using the standard EPU software along with K3 direct electron detector and a Bioquantum energy filter (Gatan Inc) with 20 eV slit. Movies were collected at 105,000× magnification resulting in a pixel size of 0.84 Å. Movies were collected at a total dose ranging from 41.7 to 58.9 e^−^/Å^2^, with 0.5 to 1.8 s exposures, and a defocus range of −0.3 to −1.3 µm.

### Image processing

All movies were processed using cryoSPARC Live and cryoSPARC ^23^. Motion correction and CTF estimation were performed using default parameters and initial particle extraction used the built-in *blob picker* with a box size of 320 pixels ^24^. Initial subsets of particles were subjected to reference free 2D classification before discreet and diverse classes were chosen to re-extract particles using template picking. Multiple rounds of classification were performed to exclude junk and non-homogenous classes. Ab-initio models were generated using a subset of these particles and C1 symmetry. The entire particle set was refined in 3D against ab-initio models without symmetry. Octahedral symmetry was imposed in subsequent rounds of refinement for ferritin while D6 symmetry was applied to PDX datasets. Per particle local CTF refinement was performed before the final round of homogenous refinement ^25^. Resolution of the final map was estimated using the gold standard at 0.143 FSC. Maps were visualized using UCSF Chimera ^26^ and have been deposited in the EMDB repository as described in the data availability statement.

## Author Contributions

JEE conceived the research. SMP performed all cell-free expression, purification and cryo-EM experiments. JEE and SMP performed all cryo-EM image analysis. SMP performed density fitting and model refinement. JEE and SMP cowrote and edited the manuscript.

## Acknowledgments/Funding

This research was partially supported by the DOE Office of Biological and Environmental Research, Biological Systems Science Division, FWP 74915. This research was performed on a project award (10.46936/intm.proj.2023.60672/60008773) from the Environmental Molecular Sciences Laboratory, a DOE Office of Science User Facility sponsored by the Biological and Environmental Research program under Contract No. DE-AC05-76RL01830.

## Data Availability Statement

The datasets for this study can be found at the PDB and EMDB repositories with the following ascension numbers that will be released with publication. *Artemia* ferritin cell-free expression lysate (EMD-49483), and variations such as expression followed by 300 kDa MWCO filtering (EMD-71215), or benzonase treatment (EMD-71217), reverse His purification (EMD-49464) and reverse His purification with 1 MDa MWCO filtration (EMD-71203). For *Arabidopsis* PDX1.2, cell-free expression lysates 60k particle (EMD-71206) and 90k particles (EMD-71207), 300 kDa MWCO filtration (EMD-71212), benzonase treatment (EMD-71209), and benzonase combined with reverse His purification (EMD-71208). The atomic model for *Artemia* ferritin derived from EMD-49464 is PDB 9P35.

