## Supplemental Information for "Cryo-EM protein structure without purification"

for

Contents:

Supplementary Figures 1–3

Supplementary Tables 1-3

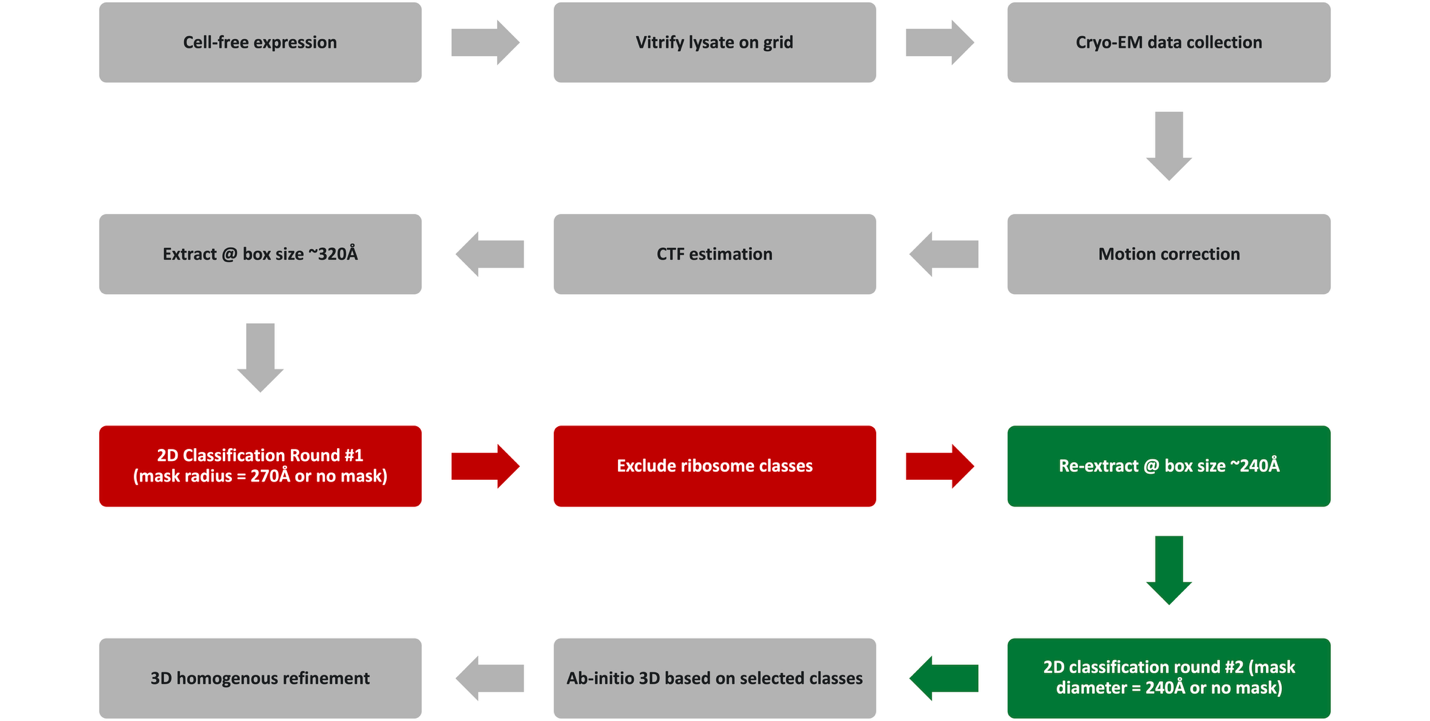
**Supplemental Figure 1. Overall workflow for image processing.** After cell-free expression of target protein finishes, standard methods for vitrification, data collection and pre-processing steps up to the first extraction are performed as normal. Then two rounds of 2D classification are used to generate the initial population of particles representing target protein. For the first round of 2D classification (red boxes) the goal is to pick everything then find and exclude the ribosomal fraction. All other non-ribosome classes are then passed to a new extraction job with a smaller box size prior to the second round of 2D classification being used to exclude junk classes and other non-target subpopulations. The selected classes of target protein then continue through either additional 2D classification jobs to triage bad particles or sent to normal ab-initio and later 3D homogenous refinement jobs.

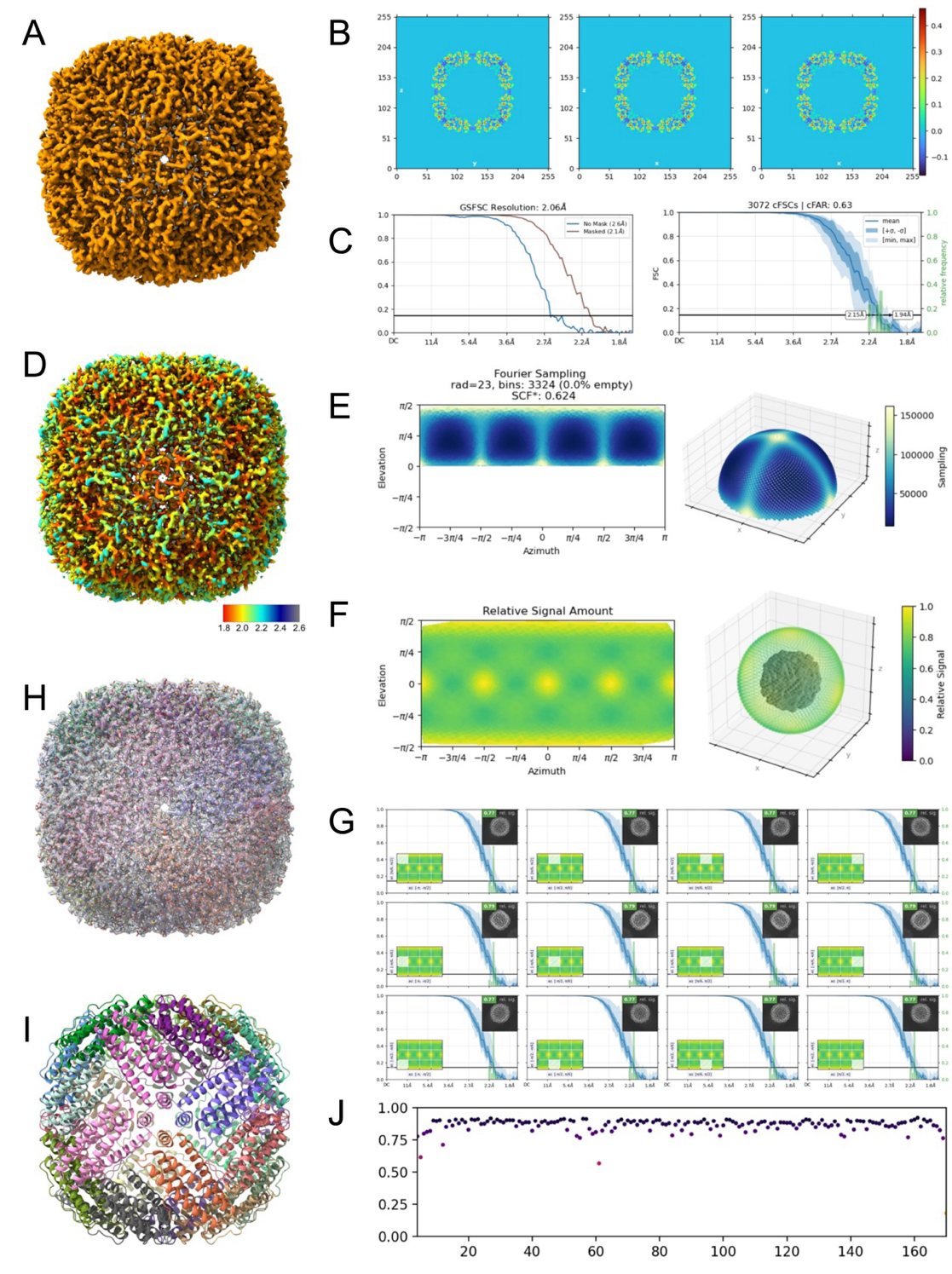

**Supplemental Figure 2. Map and model quality for *Artemia* ferritin.** A) Final map of *Artemia* ferritin from cell-free expression with reverse His purification (EMD-49464) along with corresponding Real Space Slices (B), and Gold-Standard Fourier Shell Correlation and Conical Fourier Shell Correlation plots (C). D) Local resolution colored map from (A) and corresponding Fourier Sampling plot (E), Relative Signal Amount Versus Viewing Direction (F), and Average Relative Signal Amount within Azimuth-Elevation Viewing Regions (G). H) overlaid fit between final map with refined atomic model (PDB: 9P35). I) Cartoon depiction of refined atomic model with each monomer separately colored. J) Q-score values for map and model fit demonstrating values greater than 0.8 for >96% of all amino acids.

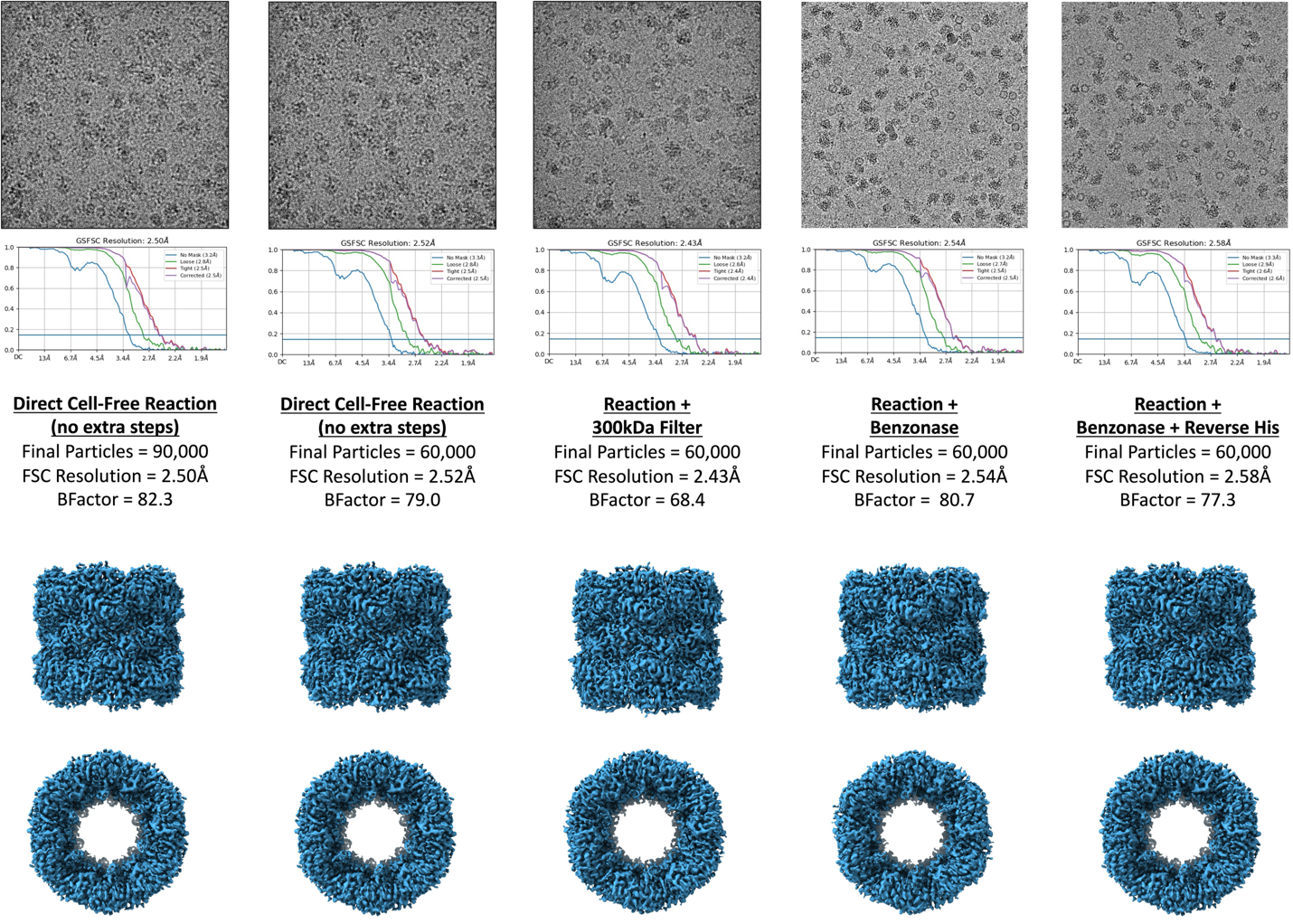

**Supplemental Figure 3. Comparison of attainable resolution for various post expression treatments of PDX1.2.** Comparative final structures of PDX1.2 without any purification, with benzonase treatment, 300 kDa MWCO spin column filtration, or Reverse His affinity tag purification using the same imaging parameters for all datasets including the number of particles in the final refinement. Note that none of the treatments significantly improved the overall resolution suggesting that PDX1.2 was likely limited by inherent structural flexibility or disorder and was already near the max attainable resolution in the crude cell-free expression lysate without purification.

**Supplement Table 1 – Dataset details for all *Artemia* ferritin samples**

| **Dataset** | ***Artemia* ferritin cell-free expression lysate** | ***Artemia* ferritin cell-free expression with 300 kDa MWCO filtering** | ***Artemia* ferritin cell-free expression with benzonase** | ***Artemia* ferritin cell-free expression with reverse His purification** | ***Artemia* ferritin cell-free expression with reverse His purification and 1 MDa MWCO Filtration** |
| --- | --- | --- | --- | --- | --- |
| EMDB ID | EMD-49483 | EMD-71215 | EMD-71217 | EMD-49464 | EMD-71203 |
| Magnification | 105 kx | 105 kx | 105 kx | 105 kx | 105 kx |
| Super resolution | No | No | No | No | No |
| Pixel size (Å) | 0.84 | 0.84 | 0.84 | 0.84 | 0.84 |
| Total dose (e-/ Å2) | 55 | 55 | 55 | 55 | 55 |
| Defocus range (µm) | 0.5-3.0 | 0.5-3.0 | 0.5-3.0 | 0.5-3.0 | 0.5-3.0 |
| Micrographs collected | 4901 | 4564 | 3580 | 3425 | 5796 |
| Micrographs utilized | 3298 | 3726 | 1319 | 2145 | 3482 |
| Total particles picked (all subclasses) | 250265 | 674815 | 252903 | 647328 | 359875 |
| Particles after first round of triage (Total particles minus ribosome sublass) * | 147859 | 160489 | 108354 | 116692 | 124547 |
| Final particle number | 90k | 90k | 90k | 90k | 90k |
| Symmetry imposed | Octahedral | Octahedral | Octahedral | Octahedral | Octahedral |
| Resolution (0.143 FSC) | 2.80 | 2.23 | 2.13 | 2.06 | 2.23 |

* Note that the traditional “initial particles / total particles picked” category is not the best metric for the current described workflow since it represents all picked particles from a complex mixture. Therefore, we created the “Particles after first round of triage” category.

**Supplement Table 2 – Model validation statistics for *Artemia* ferritin (PDB: 9P35)**

| **Composition** | |
| --- | --- |
| Chains | 24 |
| Atoms | 33816 (Hydrogens: 0) |
| Residues | Protein: 4104, Nucleotide: 0 |
| Water | 0 |
| Ligands | 0 |
| **Bonds (RMSD)** | |
| Length (Å) (# > 4s) | 0.003 (0) |
| Angles (°) (# > 4s) | 0.529 (3) |
| Molprobity score | 1.05 |
| Clashscore | 2.65 |
| **Ramachandran plot (%)** | |
| Outliers | 0.00 |
| Allowed | 0.62 |
| Favored | 99.38 |
| Rotamer outliers (%) | 0.92 |
| Cß outliers (%) | 0.00 |
| **Model vs. Data** |  |
| D FSC model (0/0.143/0.5) | 2.0/2.0/2.2 |
| CC (volume) | 0.84 |

**Supplement Table 3 – Dataset details for all *Arabidopsis* PDX1.2 samples**

| **Dataset** | PDX1.2 cell-free expression lysate 90k | PDX1.2 cell-free expression lysate 60k | PDX1.2 cell-free expression lysate with 300 kDa MWCO Filtration | PDX1.2 cell-free expression lysate with Benzonase | PDX1.2 cell-free expression lysate with Benzonase and reverse His purification |
| --- | --- | --- | --- | --- | --- |
| EMDB ID | EMD-71207 | EMD-71206 | EMD-71212 | EMD-71209 | EMD-71208 |
| Magnification | 105 kx | 105 kx | 105 kx | 105 kx | 105 kx |
| Super resolution | No | No | No | No | No |
| Pixel size (Å) | 0.84 | 0.84 | 0.84 | 0.84 | 0.84 |
| Total dose (e-/ Å2) | 55 | 55 | 55 | 55 | 55 |
| Defocus range (µm) | 0.5-3.0 | 0.5-3.0 | 0.5-3.0 | 0.5-3.0 | 0.5-3.0 |
| Micrographs collected | 4873 | 4873 | 2079 | 5215 | 1722 |
| Micrographs utilized | 3088 | 3088 | 1679 | 2415 | 1722 |
| Total particles picked (all subclasses) | 594554 | 594554 | 461933 | 573593 | 406713 |
| Particles after first round of triage (Total particles minus ribosome sublass) * | 131866 | 131866 | 183168 | 192920 | 199960 |
| Final particle number | 90k | 60k | 60k | 60k | 60k |
| Symmetry imposed | D6 | D6 | D6 | D6 | D6 |
| Resolution (0.143 FSC) | 2.50 | 2.52 | 2.43 | 2.54 | 2.58 |

* Note that the traditional “initial particles / total particles picked” category is not the best metric for the current described workflow since it represents all picked particles from a complex mixture. Therefore, we created the “Particles after first round of triage” category.
